## Supplementary material for "NCodR: A multi-class SVM classification to distinguish between non-coding RNAs in Viridiplantae"

[2]Present Address: Laboratory of Bioinformatics and Protein Engineering, International Institute of Molecular and Cell Biology in Warsaw, ul. Ks. Trojdena 4, PL-02-109 Warsaw, Poland

*Corresponding author

Corresponding author: Ranjit Prasad Bahadur

**Table S1: Performance measures of the different classifiers with 50/50 ratio of training/testing dataset**

| Classifiers | Performance measures | lncRNA | miRNA | premiRNA | rRNA | snoRNA | tRNA | snRNA | overall |
|---|---|---|---|---|---|---|---|---|---|
| k-Nearest Neighbors | Sensitivity | 0.841 | 0.283 | 0.949 | 0.989 | 0.650 | 0.933 | 0.899 | 0.917 |
| | Specificity | 0.971 | 0.999 | 0.987 | 0.966 | 0.996 | 0.991 | 0.987 | 0.986 |
| | PPV | 0.568 | 0.810 | 0.957 | 0.937 | 0.923 | 0.952 | 0.920 | 0.917 |
| | NPV | 0.993 | 0.987 | 0.985 | 0.994 | 0.976 | 0.987 | 0.983 | 0.986 |
| | FPR | 0.029 | 0.001 | 0.013 | 0.034 | 0.004 | 0.009 | 0.013 | 0.014 |
| | FNR | 0.159 | 0.717 | 0.051 | 0.011 | 0.350 | 0.067 | 0.101 | 0.083 |
| | FDR | 0.432 | 0.190 | 0.043 | 0.063 | 0.077 | 0.048 | 0.080 | 0.083 |
| | Accuracy | 0.966 | 0.986 | 0.979 | 0.974 | 0.974 | 0.981 | 0.974 | 0.976 |
| | F1 | 0.678 | 0.419 | 0.953 | 0.962 | 0.763 | 0.942 | 0.909 | 0.917 |
| Linear SVM | Sensitivity | 0.782 | 0.397 | 0.956 | 0.964 | 0.683 | 0.913 | 0.931 | 0.912 |
| | Specificity | 0.992 | 0.993 | 0.991 | 0.970 | 0.973 | 0.988 | 0.987 | 0.985 |
| | PPV | 0.824 | 0.492 | 0.968 | 0.943 | 0.641 | 0.937 | 0.923 | 0.912 |
| | NPV | 0.990 | 0.989 | 0.987 | 0.981 | 0.978 | 0.983 | 0.988 | 0.985 |
| | FPR | 0.008 | 0.007 | 0.009 | 0.030 | 0.027 | 0.012 | 0.013 | 0.015 |
| | FNR | 0.218 | 0.603 | 0.044 | 0.036 | 0.317 | 0.087 | 0.069 | 0.088 |
| | FDR | 0.176 | 0.508 | 0.032 | 0.057 | 0.359 | 0.063 | 0.077 | 0.088 |
| | Accuracy | 0.983 | 0.982 | 0.983 | 0.968 | 0.954 | 0.975 | 0.979 | 0.975 |
| | F1 | 0.802 | 0.439 | 0.962 | 0.953 | 0.661 | 0.925 | 0.927 | 0.912 |
| Decision Tree | Sensitivity | 0.785 | 0.619 | 0.963 | 0.956 | 0.795 | 0.965 | 0.928 | 0.931 |
| | Specificity | 0.992 | 0.993 | 0.989 | 0.976 | 0.986 | 0.992 | 0.988 | 0.989 |
| | PPV | 0.810 | 0.624 | 0.962 | 0.953 | 0.795 | 0.962 | 0.930 | 0.931 |
| | NPV | 0.990 | 0.993 | 0.989 | 0.978 | 0.986 | 0.993 | 0.988 | 0.989 |
| | FPR | 0.008 | 0.007 | 0.011 | 0.024 | 0.014 | 0.008 | 0.012 | 0.011 |
| | FNR | 0.215 | 0.381 | 0.037 | 0.044 | 0.205 | 0.035 | 0.072 | 0.069 |
| | FDR | 0.190 | 0.376 | 0.038 | 0.047 | 0.205 | 0.038 | 0.070 | 0.069 |

| Classifiers | Performance measures | lncRNA | miRNA | premiRNA | rRNA | snoRNA | tRNA | snRNA | overall |
|---|---|---|---|---|---|---|---|---|---|
| | Accuracy | 0.983 | 0.987 | 0.983 | 0.969 | 0.973 | 0.988 | 0.980 | 0.980 |
| | F1 | 0.797 | 0.622 | 0.963 | 0.955 | 0.795 | 0.964 | 0.929 | 0.931 |
| Random Forest | Sensitivity | 0.870 | 0.574 | 0.979 | 0.985 | 0.903 | 0.990 | 0.980 | 0.966 |
| | Specificity | 0.996 | 0.998 | 0.997 | 0.986 | 0.991 | 0.997 | 0.993 | 0.994 |
| | PPV | 0.899 | 0.835 | 0.991 | 0.973 | 0.881 | 0.984 | 0.960 | 0.966 |
| | NPV | 0.994 | 0.992 | 0.994 | 0.992 | 0.993 | 0.998 | 0.997 | 0.994 |
| | FPR | 0.004 | 0.002 | 0.003 | 0.014 | 0.009 | 0.003 | 0.007 | 0.006 |
| | FNR | 0.130 | 0.426 | 0.021 | 0.015 | 0.097 | 0.010 | 0.020 | 0.034 |
| | FDR | 0.101 | 0.165 | 0.009 | 0.027 | 0.119 | 0.016 | 0.040 | 0.034 |
| | Accuracy | 0.990 | 0.990 | 0.993 | 0.986 | 0.986 | 0.996 | 0.991 | 0.990 |
| | F1 | 0.884 | 0.681 | 0.985 | 0.979 | 0.892 | 0.987 | 0.970 | 0.966 |
| AdaBoost | Sensitivity | 0.268 | 0.625 | 0.936 | 0.787 | 0.398 | 0.791 | 0.516 | 0.732 |
| | Specificity | 0.988 | 0.979 | 0.971 | 0.935 | 0.922 | 0.961 | 0.925 | 0.955 |
| | PPV | 0.504 | 0.347 | 0.903 | 0.861 | 0.262 | 0.803 | 0.535 | 0.732 |
| | NPV | 0.968 | 0.993 | 0.981 | 0.896 | 0.956 | 0.958 | 0.92 | 0.955 |
| | FPR | 0.012 | 0.021 | 0.029 | 0.065 | 0.078 | 0.039 | 0.075 | 0.045 |
| | FNR | 0.732 | 0.375 | 0.064 | 0.213 | 0.602 | 0.209 | 0.484 | 0.268 |
| | FDR | 0.496 | 0.653 | 0.097 | 0.139 | 0.738 | 0.197 | 0.465 | 0.268 |
| | Accuracy | 0.957 | 0.972 | 0.963 | 0.885 | 0.887 | 0.933 | 0.866 | 0.923 |
| | F1 | 0.35 | 0.446 | 0.919 | 0.822 | 0.316 | 0.797 | 0.525 | 0.732 |
| Naive Bayes | Sensitivity | 0.808 | 0.917 | 0.827 | 0.786 | 0.164 | 0.880 | 0.914 | 0.792 |
| | Specificity | 0.963 | 0.974 | 0.993 | 0.974 | 0.977 | 0.930 | 0.948 | 0.965 |
| | PPV | 0.496 | 0.392 | 0.973 | 0.939 | 0.333 | 0.716 | 0.745 | 0.792 |
| | NPV | 0.991 | 0.998 | 0.952 | 0.899 | 0.943 | 0.975 | 0.985 | 0.965 |
| | FPR | 0.037 | 0.026 | 0.007 | 0.026 | 0.023 | 0.070 | 0.052 | 0.035 |
| | FNR | 0.192 | 0.083 | 0.173 | 0.214 | 0.836 | 0.120 | 0.086 | 0.208 |
| | FDR | 0.504 | 0.608 | 0.027 | 0.061 | 0.667 | 0.284 | 0.255 | 0.208 |

| Classifiers | Performance measures | lncRNA | miRNA | premiRNA | rRNA | snoRNA | tRNA | snRNA | overall |
|---|---|---|---|---|---|---|---|---|---|
| | Accuracy | 0.956 | 0.973 | 0.956 | 0.911 | 0.924 | 0.922 | 0.943 | 0.941 |
| | F1 | 0.615 | 0.549 | 0.894 | 0.856 | 0.219 | 0.790 | 0.821 | 0.792 |
| QDA | Sensitivity | 0.902 | 0.938 | 0.878 | 0.577 | 0.180 | 0.923 | 0.946 | 0.750 |
| | Specificity | 0.883 | 0.975 | 0.982 | 0.996 | 0.982 | 0.948 | 0.957 | 0.958 |
| | PPV | 0.258 | 0.409 | 0.935 | 0.986 | 0.407 | 0.781 | 0.788 | 0.750 |
| | NPV | 0.995 | 0.999 | 0.965 | 0.822 | 0.945 | 0.984 | 0.991 | 0.958 |
| | FPR | 0.117 | 0.025 | 0.018 | 0.004 | 0.018 | 0.052 | 0.043 | 0.042 |
| | FNR | 0.098 | 0.062 | 0.122 | 0.423 | 0.820 | 0.077 | 0.054 | 0.250 |
| | FDR | 0.742 | 0.591 | 0.065 | 0.014 | 0.593 | 0.219 | 0.212 | 0.250 |
| | Accuracy | 0.884 | 0.975 | 0.959 | 0.855 | 0.929 | 0.944 | 0.956 | 0.929 |
| | F1 | 0.401 | 0.569 | 0.906 | 0.728 | 0.250 | 0.846 | 0.86 | 0.75 |
| RBF SVM | Sensitivity | 0.875 | 0.442 | 0.977 | 0.986 | 0.926 | 0.982 | 0.979 | 0.964 |
| | Specificity | 0.994 | 0.998 | 0.996 | 0.991 | 0.987 | 0.997 | 0.995 | 0.994 |
| | PPV | 0.874 | 0.799 | 0.986 | 0.982 | 0.833 | 0.984 | 0.969 | 0.964 |
| | NPV | 0.994 | 0.990 | 0.993 | 0.993 | 0.995 | 0.996 | 0.996 | 0.994 |
| | FPR | 0.006 | 0.002 | 0.004 | 0.009 | 0.013 | 0.003 | 0.005 | 0.006 |
| | FNR | 0.125 | 0.558 | 0.023 | 0.014 | 0.074 | 0.018 | 0.021 | 0.036 |
| | FDR | 0.126 | 0.201 | 0.014 | 0.018 | 0.167 | 0.016 | 0.031 | 0.036 |
| | Accuracy | 0.989 | 0.988 | 0.992 | 0.989 | 0.983 | 0.994 | 0.992 | 0.990 |
| | F1 | 0.874 | 0.569 | 0.982 | 0.984 | 0.877 | 0.983 | 0.974 | 0.964 |
| Meta-classifier | Sensitivity | 0.886 | 0.821 | 0.963 | 0.974 | 0.755 | 0.977 | 0.977 | 0.952 |
| | Specificity | 0.992 | 0.990 | 0.997 | 0.989 | 0.993 | 0.988 | 0.994 | 0.992 |
| | PPV | 0.825 | 0.602 | 0.990 | 0.978 | 0.889 | 0.932 | 0.972 | 0.952 |
| | NPV | 0.995 | 0.997 | 0.989 | 0.987 | 0.983 | 0.996 | 0.995 | 0.992 |
| | FPR | 0.008 | 0.010 | 0.003 | 0.011 | 0.007 | 0.012 | 0.006 | 0.008 |
| | FNR | 0.114 | 0.179 | 0.037 | 0.026 | 0.245 | 0.023 | 0.023 | 0.048 |

| Classifiers | Performance measures | lncRNA | miRNA | premiRNA | rRNA | snoRNA | tRNA | snRNA | overall |
|---|---|---|---|---|---|---|---|---|---|
| | FDR | 0.175 | 0.398 | 0.010 | 0.022 | 0.111 | 0.068 | 0.028 | 0.048 |
| | Accuracy | 0.987 | 0.987 | 0.989 | 0.984 | 0.978 | 0.987 | 0.991 | 0.986 |
| | F1 | 0.854 | 0.695 | 0.976 | 0.976 | 0.816 | 0.954 | 0.975 | 0.952 |

**Table S2: Performance measures of the different classifiers with 80/20 ratio of training/testing dataset**

| Classifiers | Performance measures | lncRNA | miRNA | premiRNA | rRNA | snoRNA | tRNA | snRNA | overall |
|---|---|---|---|---|---|---|---|---|---|
| k-Nearest Neighbors | Sensitivity | 0.855 | 0.316 | 0.958 | 0.988 | 0.679 | 0.917 | 0.946 | 0.927 |
| | Specificity | 0.975 | 0.999 | 0.989 | 0.972 | 0.997 | 0.989 | 0.991 | 0.988 |
| | PPV | 0.603 | 0.797 | 0.961 | 0.948 | 0.936 | 0.931 | 0.954 | 0.927 |
| | NPV | 0.993 | 0.988 | 0.988 | 0.994 | 0.978 | 0.986 | 0.989 | 0.988 |
| | FPR | 0.025 | 0.001 | 0.011 | 0.028 | 0.003 | 0.011 | 0.009 | 0.012 |
| | FNR | 0.145 | 0.684 | 0.042 | 0.012 | 0.321 | 0.083 | 0.054 | 0.073 |
| | FDR | 0.397 | 0.203 | 0.039 | 0.052 | 0.064 | 0.069 | 0.046 | 0.073 |
| | Accuracy | 0.970 | 0.987 | 0.982 | 0.978 | 0.976 | 0.978 | 0.983 | 0.979 |
| | F1 | 0.707 | 0.453 | 0.960 | 0.968 | 0.787 | 0.924 | 0.950 | 0.927 |
| Linear SVM | Sensitivity | 0.775 | 0.404 | 0.955 | 0.964 | 0.677 | 0.932 | 0.909 | 0.912 |
| | Specificity | 0.992 | 0.993 | 0.991 | 0.969 | 0.973 | 0.987 | 0.988 | 0.985 |
| | PPV | 0.822 | 0.501 | 0.967 | 0.941 | 0.639 | 0.923 | 0.936 | 0.912 |
| | NPV | 0.990 | 0.990 | 0.987 | 0.981 | 0.977 | 0.989 | 0.982 | 0.985 |
| | FPR | 0.008 | 0.007 | 0.009 | 0.031 | 0.027 | 0.013 | 0.012 | 0.015 |
| | FNR | 0.225 | 0.596 | 0.045 | 0.036 | 0.323 | 0.068 | 0.091 | 0.088 |
| | FDR | 0.178 | 0.499 | 0.033 | 0.059 | 0.361 | 0.077 | 0.064 | 0.088 |
| | Accuracy | 0.983 | 0.983 | 0.983 | 0.967 | 0.954 | 0.979 | 0.974 | 0.975 |

| Classifiers | Performance measures | lncRNA | miRNA | premiRNA | rRNA | snoRNA | tRNA | snRNA | overall |
|---|---|---|---|---|---|---|---|---|---|
| | F1 | 0.798 | 0.447 | 0.961 | 0.952 | 0.657 | 0.928 | 0.922 | 0.912 |
| Decision Tree | Sensitivity | 0.786 | 0.643 | 0.967 | 0.960 | 0.801 | 0.936 | 0.968 | 0.936 |
| | Specificity | 0.992 | 0.993 | 0.990 | 0.978 | 0.987 | 0.989 | 0.993 | 0.989 |
| | PPV | 0.814 | 0.626 | 0.965 | 0.957 | 0.809 | 0.936 | 0.967 | 0.936 |
| | NPV | 0.990 | 0.994 | 0.990 | 0.979 | 0.986 | 0.989 | 0.994 | 0.989 |
| | FPR | 0.008 | 0.007 | 0.010 | 0.022 | 0.013 | 0.011 | 0.007 | 0.011 |
| | FNR | 0.214 | 0.357 | 0.033 | 0.040 | 0.199 | 0.064 | 0.032 | 0.064 |
| | FDR | 0.186 | 0.374 | 0.035 | 0.043 | 0.191 | 0.064 | 0.033 | 0.064 |
| | Accuracy | 0.983 | 0.987 | 0.985 | 0.972 | 0.975 | 0.982 | 0.989 | 0.982 |
| | F1 | 0.800 | 0.634 | 0.966 | 0.959 | 0.805 | 0.936 | 0.968 | 0.936 |
| Random Forest | Sensitivity | 0.889 | 0.620 | 0.982 | 0.986 | 0.909 | 0.983 | 0.991 | 0.970 |
| | Specificity | 0.996 | 0.998 | 0.998 | 0.988 | 0.993 | 0.994 | 0.997 | 0.995 |
| | PPV | 0.905 | 0.840 | 0.992 | 0.976 | 0.895 | 0.965 | 0.986 | 0.970 |
| | NPV | 0.995 | 0.993 | 0.995 | 0.993 | 0.994 | 0.997 | 0.998 | 0.995 |
| | FPR | 0.004 | 0.002 | 0.002 | 0.012 | 0.007 | 0.006 | 0.003 | 0.005 |
| | FNR | 0.111 | 0.380 | 0.018 | 0.014 | 0.091 | 0.017 | 0.009 | 0.030 |
| | FDR | 0.095 | 0.160 | 0.008 | 0.024 | 0.105 | 0.035 | 0.014 | 0.030 |
| | Accuracy | 0.991 | 0.991 | 0.994 | 0.987 | 0.987 | 0.992 | 0.996 | 0.991 |
| | F1 | 0.897 | 0.713 | 0.987 | 0.981 | 0.902 | 0.974 | 0.988 | 0.970 |
| AdaBoost | Sensitivity | 0.372 | 0.532 | 0.895 | 0.862 | 0.414 | 0.025 | 0.763 | 0.677 |
| | Specificity | 0.981 | 0.980 | 0.985 | 0.823 | 0.890 | 0.976 | 0.960 | 0.946 |
| | PPV | 0.464 | 0.322 | 0.947 | 0.714 | 0.207 | 0.149 | 0.792 | 0.677 |
| | NPV | 0.972 | 0.992 | 0.970 | 0.921 | 0.956 | 0.857 | 0.953 | 0.946 |
| | FPR | 0.019 | 0.020 | 0.015 | 0.177 | 0.110 | 0.024 | 0.040 | 0.054 |
| | FNR | 0.628 | 0.468 | 0.105 | 0.138 | 0.586 | 0.975 | 0.237 | 0.323 |

| Classifiers | Performance measures | lncRNA | miRNA | premiRNA | rRNA | snoRNA | tRNA | snRNA | overall |
|---|---|---|---|---|---|---|---|---|---|
| | FDR | 0.536 | 0.678 | 0.053 | 0.286 | 0.793 | 0.851 | 0.208 | 0.323 |
| | Accuracy | 0.955 | 0.973 | 0.965 | 0.836 | 0.859 | 0.840 | 0.927 | 0.908 |
| | F1 | 0.413 | 0.401 | 0.921 | 0.781 | 0.276 | 0.042 | 0.777 | 0.677 |
| Naive Bayes | Sensitivity | 0.811 | 0.926 | 0.827 | 0.784 | 0.154 | 0.913 | 0.883 | 0.791 |
| | Specificity | 0.963 | 0.974 | 0.993 | 0.974 | 0.977 | 0.948 | 0.929 | 0.965 |
| | PPV | 0.495 | 0.384 | 0.972 | 0.939 | 0.321 | 0.746 | 0.712 | 0.791 |
| | NPV | 0.991 | 0.999 | 0.952 | 0.898 | 0.943 | 0.985 | 0.975 | 0.965 |
| | FPR | 0.037 | 0.026 | 0.007 | 0.026 | 0.023 | 0.052 | 0.071 | 0.035 |
| | FNR | 0.189 | 0.074 | 0.173 | 0.216 | 0.846 | 0.087 | 0.117 | 0.209 |
| | FDR | 0.505 | 0.616 | 0.028 | 0.061 | 0.679 | 0.254 | 0.288 | 0.209 |
| | Accuracy | 0.956 | 0.973 | 0.956 | 0.910 | 0.924 | 0.943 | 0.921 | 0.940 |
| | F1 | 0.615 | 0.543 | 0.894 | 0.855 | 0.208 | 0.821 | 0.789 | 0.791 |
| QDA | Sensitivity | 0.899 | 0.965 | 0.876 | 0.583 | 0.153 | 0.946 | 0.923 | 0.750 |
| | Specificity | 0.885 | 0.974 | 0.982 | 0.996 | 0.981 | 0.958 | 0.947 | 0.958 |
| | PPV | 0.259 | 0.394 | 0.934 | 0.986 | 0.365 | 0.790 | 0.778 | 0.750 |
| | NPV | 0.995 | 0.999 | 0.964 | 0.823 | 0.943 | 0.991 | 0.984 | 0.958 |
| | FPR | 0.115 | 0.026 | 0.018 | 0.004 | 0.019 | 0.042 | 0.053 | 0.042 |
| | FNR | 0.101 | 0.035 | 0.124 | 0.417 | 0.847 | 0.054 | 0.077 | 0.250 |
| | FDR | 0.741 | 0.606 | 0.066 | 0.014 | 0.635 | 0.210 | 0.222 | 0.250 |
| | Accuracy | 0.885 | 0.974 | 0.958 | 0.856 | 0.927 | 0.956 | 0.943 | 0.929 |
| | F1 | 0.402 | 0.559 | 0.904 | 0.733 | 0.216 | 0.861 | 0.844 | 0.750 |
| RBF SVM | Sensitivity | 0.808 | 0.044 | 0.966 | 0.981 | 0.928 | 0.965 | 0.969 | 0.947 |
| | Specificity | 0.993 | 1.000 | 0.995 | 0.985 | 0.978 | 0.992 | 0.994 | 0.991 |
| | PPV | 0.841 | 0.798 | 0.982 | 0.972 | 0.742 | 0.950 | 0.972 | 0.947 |
| | NPV | 0.991 | 0.984 | 0.990 | 0.990 | 0.995 | 0.994 | 0.994 | 0.991 |

| Classifiers | Performance measures | lncRNA | miRNA | premiRNA | rRNA | snoRNA | tRNA | snRNA | overall |
|---|---|---|---|---|---|---|---|---|---|
| | FPR | 0.007 | 0.000 | 0.005 | 0.015 | 0.022 | 0.008 | 0.006 | 0.009 |
| | FNR | 0.192 | 0.956 | 0.034 | 0.019 | 0.072 | 0.035 | 0.031 | 0.053 |
| | FDR | 0.159 | 0.202 | 0.018 | 0.028 | 0.258 | 0.050 | 0.028 | 0.053 |
| | Accuracy | 0.985 | 0.983 | 0.988 | 0.984 | 0.974 | 0.988 | 0.990 | 0.985 |
| | F1 | 0.824 | 0.083 | 0.974 | 0.976 | 0.825 | 0.958 | 0.970 | 0.947 |
| Meta-classifier | Sensitivity | 0.892 | 0.848 | 0.968 | 0.975 | 0.767 | 0.981 | 0.981 | 0.956 |
| | Specificity | 0.992 | 0.990 | 0.997 | 0.990 | 0.995 | 0.989 | 0.995 | 0.993 |
| | PPV | 0.836 | 0.608 | 0.991 | 0.980 | 0.908 | 0.939 | 0.975 | 0.956 |
| | NPV | 0.995 | 0.997 | 0.991 | 0.987 | 0.984 | 0.997 | 0.996 | 0.993 |
| | FPR | 0.008 | 0.010 | 0.003 | 0.010 | 0.005 | 0.011 | 0.005 | 0.007 |
| | FNR | 0.108 | 0.152 | 0.032 | 0.025 | 0.233 | 0.019 | 0.019 | 0.044 |
| | FDR | 0.164 | 0.392 | 0.009 | 0.020 | 0.092 | 0.061 | 0.025 | 0.044 |
| | Accuracy | 0.988 | 0.988 | 0.991 | 0.985 | 0.980 | 0.988 | 0.993 | 0.987 |
| | F1 | 0.863 | 0.708 | 0.979 | 0.978 | 0.832 | 0.959 | 0.978 | 0.956 |

**Table S3: Performance measures of the different classifiers with 99.5/0.5 ratio of training/testing dataset**

| Classifiers | Performance measures | lncRNA | miRNA | premiRNA | rRNA | snoRNA | tRNA | snRNA | overall |
|---|---|---|---|---|---|---|---|---|---|
| k-Nearest Neighbors | Sensitivity | 0.867 | 0.333 | 0.960 | 0.989 | 0.679 | 0.925 | 0.951 | 0.930 |
| | Specificity | 0.977 | 0.998 | 0.989 | 0.974 | 0.996 | 0.989 | 0.991 | 0.988 |
| | PPV | 0.624 | 0.785 | 0.963 | 0.951 | 0.927 | 0.933 | 0.954 | 0.930 |
| | NPV | 0.994 | 0.988 | 0.988 | 0.994 | 0.978 | 0.988 | 0.990 | 0.988 |
| | FPR | 0.023 | 0.002 | 0.011 | 0.026 | 0.004 | 0.011 | 0.009 | 0.012 |
| | FNR | 0.133 | 0.667 | 0.040 | 0.011 | 0.321 | 0.075 | 0.049 | 0.070 |

| Classifiers | Performance measures | lncRNA | miRNA | premiRNA | rRNA | snoRNA | tRNA | snRNA | overall |
|---|---|---|---|---|---|---|---|---|---|
| | FDR | 0.376 | 0.215 | 0.037 | 0.049 | 0.073 | 0.067 | 0.046 | 0.070 |
| | Accuracy | 0.972 | 0.986 | 0.983 | 0.979 | 0.976 | 0.980 | 0.984 | 0.980 |
| | F1 | 0.726 | 0.467 | 0.962 | 0.970 | 0.784 | 0.929 | 0.953 | 0.930 |
| Linear SVM | Sensitivity | 0.785 | 0.369 | 0.953 | 0.962 | 0.698 | 0.922 | 0.911 | 0.911 |
| | Specificity | 0.992 | 0.994 | 0.991 | 0.969 | 0.971 | 0.987 | 0.987 | 0.985 |
| | PPV | 0.810 | 0.513 | 0.970 | 0.941 | 0.628 | 0.924 | 0.936 | 0.911 |
| | NPV | 0.990 | 0.989 | 0.986 | 0.981 | 0.979 | 0.987 | 0.982 | 0.985 |
| | FPR | 0.008 | 0.006 | 0.009 | 0.031 | 0.029 | 0.013 | 0.013 | 0.015 |
| | FNR | 0.215 | 0.631 | 0.047 | 0.038 | 0.302 | 0.078 | 0.089 | 0.089 |
| | FDR | 0.190 | 0.487 | 0.030 | 0.059 | 0.372 | 0.076 | 0.064 | 0.089 |
| | Accuracy | 0.983 | 0.982 | 0.983 | 0.967 | 0.954 | 0.978 | 0.974 | 0.974 |
| | F1 | 0.797 | 0.429 | 0.962 | 0.951 | 0.661 | 0.923 | 0.923 | 0.911 |
| Decision Tree | Sensitivity | 0.812 | 0.627 | 0.972 | 0.960 | 0.817 | 0.933 | 0.972 | 0.939 |
| | Specificity | 0.992 | 0.994 | 0.991 | 0.979 | 0.987 | 0.990 | 0.993 | 0.990 |
| | PPV | 0.827 | 0.655 | 0.971 | 0.959 | 0.809 | 0.937 | 0.967 | 0.939 |
| | NPV | 0.992 | 0.993 | 0.992 | 0.980 | 0.987 | 0.989 | 0.994 | 0.990 |
| | FPR | 0.008 | 0.006 | 0.009 | 0.021 | 0.013 | 0.010 | 0.007 | 0.010 |
| | FNR | 0.188 | 0.373 | 0.028 | 0.040 | 0.183 | 0.067 | 0.028 | 0.061 |
| | FDR | 0.173 | 0.345 | 0.029 | 0.041 | 0.191 | 0.063 | 0.033 | 0.061 |
| | Accuracy | 0.985 | 0.987 | 0.987 | 0.973 | 0.976 | 0.982 | 0.990 | 0.983 |
| | F1 | 0.819 | 0.641 | 0.971 | 0.959 | 0.813 | 0.935 | 0.970 | 0.939 |
| Random Forest | Sensitivity | 0.888 | 0.625 | 0.982 | 0.986 | 0.902 | 0.981 | 0.992 | 0.969 |
| | Specificity | 0.996 | 0.997 | 0.997 | 0.988 | 0.992 | 0.994 | 0.997 | 0.995 |
| | PPV | 0.905 | 0.815 | 0.991 | 0.976 | 0.893 | 0.967 | 0.987 | 0.969 |
| | NPV | 0.995 | 0.993 | 0.995 | 0.993 | 0.993 | 0.997 | 0.998 | 0.995 |

| Classifiers | Performance measures | lncRNA | miRNA | premiRNA | rRNA | snoRNA | tRNA | snRNA | overall |
|---|---|---|---|---|---|---|---|---|---|
| | FPR | 0.004 | 0.003 | 0.003 | 0.012 | 0.008 | 0.006 | 0.003 | 0.005 |
| | FNR | 0.113 | 0.375 | 0.018 | 0.014 | 0.098 | 0.019 | 0.008 | 0.031 |
| | FDR | 0.095 | 0.185 | 0.009 | 0.024 | 0.107 | 0.033 | 0.013 | 0.031 |
| | Accuracy | 0.991 | 0.991 | 0.994 | 0.987 | 0.987 | 0.992 | 0.996 | 0.991 |
| | F1 | 0.896 | 0.707 | 0.987 | 0.981 | 0.897 | 0.973 | 0.989 | 0.969 |
| AdaBoost | Sensitivity | 0.903 | 0.000 | 0.779 | 0.644 | 0.550 | 0.459 | 0.345 | 0.590 |
| | Specificity | 0.893 | 1.000 | 0.984 | 0.892 | 0.888 | 0.936 | 0.923 | 0.932 |
| | PPV | 0.273 | nan | 0.934 | 0.753 | 0.254 | 0.541 | 0.480 | 0.590 |
| | NPV | 0.995 | 0.982 | 0.939 | 0.831 | 0.966 | 0.913 | 0.872 | 0.932 |
| | FPR | 0.107 | 0.000 | 0.016 | 0.108 | 0.112 | 0.064 | 0.077 | 0.068 |
| | FNR | 0.097 | 1.000 | 0.221 | 0.356 | 0.450 | 0.541 | 0.655 | 0.410 |
| | FDR | 0.727 | nan | 0.066 | 0.247 | 0.746 | 0.459 | 0.520 | 0.410 |
| | Accuracy | 0.894 | 0.982 | 0.938 | 0.808 | 0.866 | 0.869 | 0.824 | 0.883 |
| | F1 | 0.420 | 0.000 | 0.849 | 0.694 | 0.347 | 0.496 | 0.401 | 0.590 |
| Naive Bayes | Sensitivity | 0.826 | 0.913 | 0.831 | 0.781 | 0.149 | 0.912 | 0.882 | 0.791 |
| | Specificity | 0.962 | 0.974 | 0.992 | 0.973 | 0.977 | 0.949 | 0.930 | 0.965 |
| | PPV | 0.492 | 0.394 | 0.969 | 0.937 | 0.308 | 0.747 | 0.721 | 0.791 |
| | NPV | 0.992 | 0.998 | 0.953 | 0.897 | 0.943 | 0.985 | 0.975 | 0.965 |
| | FPR | 0.038 | 0.026 | 0.008 | 0.027 | 0.023 | 0.051 | 0.070 | 0.035 |
| | FNR | 0.174 | 0.087 | 0.169 | 0.219 | 0.851 | 0.088 | 0.118 | 0.209 |
| | FDR | 0.508 | 0.606 | 0.031 | 0.063 | 0.692 | 0.253 | 0.279 | 0.209 |
| | Accuracy | 0.956 | 0.973 | 0.956 | 0.909 | 0.923 | 0.944 | 0.921 | 0.940 |
| | F1 | 0.617 | 0.550 | 0.895 | 0.852 | 0.200 | 0.821 | 0.793 | 0.791 |
| QDA | Sensitivity | 0.897 | 0.958 | 0.881 | 0.579 | 0.146 | 0.941 | 0.922 | 0.749 |
| | Specificity | 0.884 | 0.974 | 0.981 | 0.996 | 0.981 | 0.958 | 0.947 | 0.958 |

| Classifiers | Performance measures | lncRNA | miRNA | premiRNA | rRNA | snoRNA | tRNA | snRNA | overall |
|---|---|---|---|---|---|---|---|---|---|
| | PPV | 0.256 | 0.404 | 0.932 | 0.986 | 0.353 | 0.787 | 0.781 | 0.749 |
| | NPV | 0.995 | 0.999 | 0.966 | 0.823 | 0.943 | 0.990 | 0.983 | 0.958 |
| | FPR | 0.116 | 0.026 | 0.019 | 0.004 | 0.019 | 0.042 | 0.053 | 0.042 |
| | FNR | 0.103 | 0.042 | 0.119 | 0.421 | 0.854 | 0.059 | 0.078 | 0.251 |
| | FDR | 0.744 | 0.596 | 0.068 | 0.014 | 0.647 | 0.213 | 0.219 | 0.251 |
| | Accuracy | 0.885 | 0.974 | 0.959 | 0.855 | 0.927 | 0.956 | 0.943 | 0.928 |
| | F1 | 0.398 | 0.568 | 0.906 | 0.729 | 0.207 | 0.857 | 0.846 | 0.749 |
| RBF SVM | Sensitivity | 0.814 | 0.049 | 0.965 | 0.981 | 0.932 | 0.964 | 0.969 | 0.946 |
| | Specificity | 0.992 | 1.000 | 0.995 | 0.985 | 0.977 | 0.991 | 0.995 | 0.991 |
| | PPV | 0.828 | 0.793 | 0.982 | 0.971 | 0.742 | 0.949 | 0.974 | 0.946 |
| | NPV | 0.992 | 0.983 | 0.990 | 0.990 | 0.995 | 0.994 | 0.994 | 0.991 |
| | FPR | 0.008 | 0.000 | 0.005 | 0.015 | 0.023 | 0.009 | 0.005 | 0.009 |
| | FNR | 0.186 | 0.951 | 0.035 | 0.019 | 0.068 | 0.036 | 0.031 | 0.054 |
| | FDR | 0.172 | 0.207 | 0.018 | 0.029 | 0.258 | 0.051 | 0.026 | 0.054 |
| | Accuracy | 0.985 | 0.983 | 0.988 | 0.984 | 0.975 | 0.988 | 0.990 | 0.985 |
| | F1 | 0.821 | 0.092 | 0.974 | 0.976 | 0.826 | 0.956 | 0.972 | 0.946 |
| Meta-classifier | Sensitivity | 0.889 | 0.835 | 0.970 | 0.975 | 0.772 | 0.977 | 0.980 | 0.956 |
| | Specificity | 0.992 | 0.990 | 0.997 | 0.990 | 0.994 | 0.989 | 0.995 | 0.993 |
| | PPV | 0.837 | 0.613 | 0.991 | 0.981 | 0.906 | 0.937 | 0.974 | 0.956 |
| | NPV | 0.995 | 0.997 | 0.991 | 0.987 | 0.984 | 0.996 | 0.996 | 0.993 |
| | FPR | 0.008 | 0.010 | 0.003 | 0.010 | 0.006 | 0.011 | 0.005 | 0.007 |
| | FNR | 0.111 | 0.165 | 0.030 | 0.025 | 0.228 | 0.023 | 0.020 | 0.044 |
| | FDR | 0.163 | 0.387 | 0.009 | 0.019 | 0.094 | 0.063 | 0.026 | 0.044 |
| | Accuracy | 0.988 | 0.988 | 0.991 | 0.985 | 0.980 | 0.988 | 0.992 | 0.987 |
| | F1 | 0.862 | 0.707 | 0.980 | 0.978 | 0.834 | 0.957 | 0.977 | 0.956 |

**Table S4: Performance measures of the different classifiers**

| SVM-RBF | | | |
|---|---|---|---|
| ncRNA Classes | precision | recall | f1-score |
| lncRNA | 0.87 | 0.89 | 0.88 |
| miRNA | 0.85 | 0.54 | 0.66 |
| premiRNA | 0.98 | 0.98 | 0.98 |
| rRNA | 0.97 | 0.98 | 0.97 |
| snoRNA | 0.99 | 0.99 | 0.99 |
| snRNA | 0.86 | 0.94 | 0.90 |
| tRNA | 0.98 | 0.97 | 0.98 |
| Random Forest | | | |
| lncRNA | 0.89 | 0.92 | 0.91 |
| miRNA | 0.70 | 0.61 | 0.65 |
| premiRNA | 0.98 | 0.98 | 0.98 |
| rRNA | 0.95 | 0.98 | 0.97 |
| snoRNA | 0.99 | 0.98 | 0.99 |
| snRNA | 0.84 | 0.87 | 0.85 |
| tRNA | 0.99 | 0.98 | 0.98 |

**Figure S1. Confusion matrix for SVM-RBF model trained and tested on 99.5 % and 0.5% of the entire dataset respectively.**

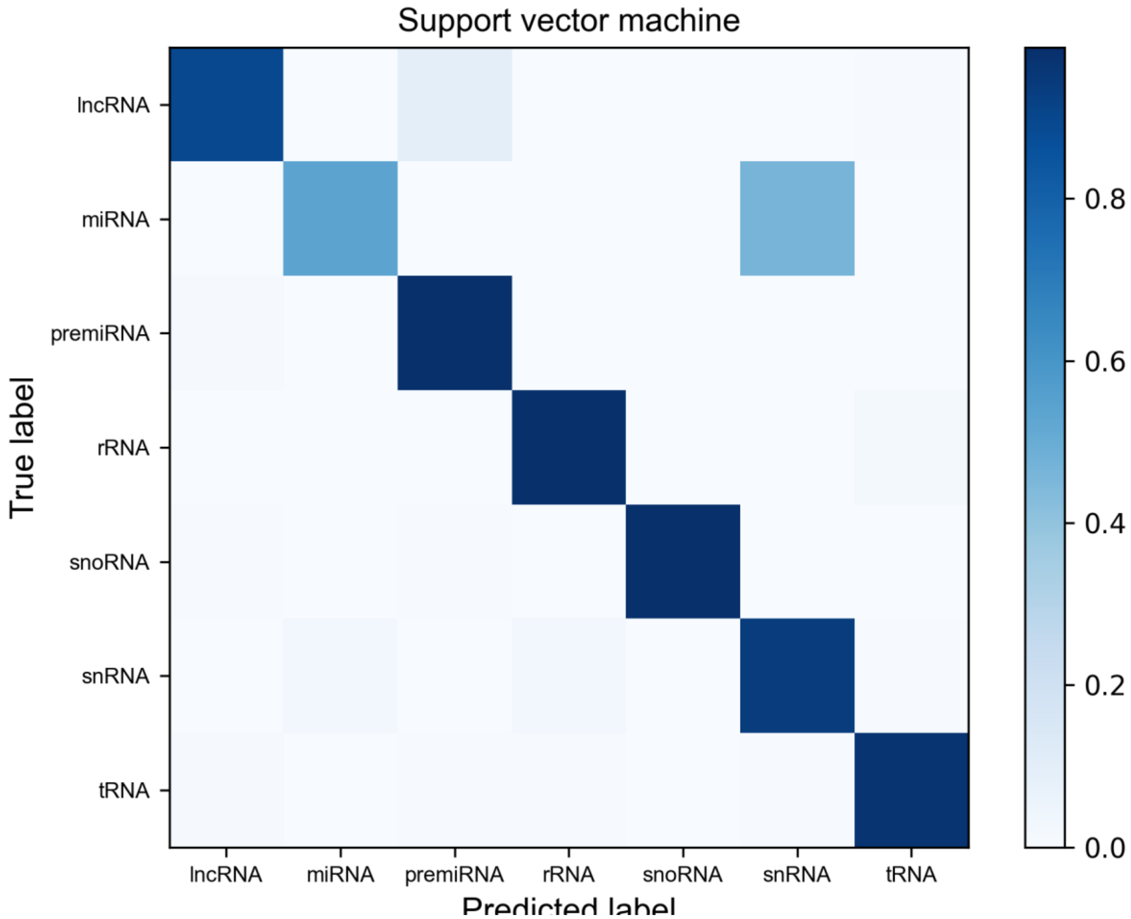
